# Predictive Metagenomic Analysis of Autoimmune Disease Identifies Robust Autoimmunity and Disease Specific Microbial Signatures

**DOI:** 10.1101/779967

**Authors:** Angelina Volkova, Kelly V. Ruggles

## Abstract

Within the last decade, numerous studies have demonstrated changes in the gut microbiome associated with specific autoimmune diseases. Due to differences in study design, data quality control, analysis and statistical methods, many results of these studies are inconsistent and incomparable. To better understand the relationship between the intestinal microbiome and autoimmunity, we have completed a comprehensive re-analysis of 42 studies focusing on the gut microbiome in twelve autoimmune diseases to identify a microbial signature predictive of multiple sclerosis (MS), inflammatory bowel disease (IBD), rheumatoid arthritis (RA) and general autoimmune disease using both 16S rRNA sequencing data and shotgun metagenomics data. To do this, we used four machine learning algorithms, random forest, eXtreme Gradient Boosting (XGBoost), ridge regression and support vector machine with radial kernel and recursive feature elimination to rank disease predictive taxa comparing disease vs. healthy participants and pairwise comparisons of each disease. Comparing the performance of these models, we found XGBoost and random forest, tree-based methods capable of handling sparse multidimensional data, to consistently produce the best results. Through this modeling, we identified a number of taxa consistently identified as dysregulated in a general autoimmune disease model including *Odoribacter, Lachnospiraceae Clostridium* and *Mogibacteriaceae* implicating all as potential factors connecting the gut microbiome and to autoimmune response. Further, we computed pairwise comparison models to identify disease specific taxa signatures highlighting a role for *Peptostreptococcaceae* and *Ruminococcaceae Gemmiger* in IBD and *Akkermansia, Butyricicoccus and Mogibacteriaceae* in MS. We then connected a subset of these taxa with potential metabolic alterations based on metagenomic/metabolomic correlation analysis, identifying 250 metabolites associated with autoimmunity-predictive taxa.

## INTRODUCTION

The human intestine is colonized by millions of microbes, which have been shown to be involved in metabolism^1^, immunity^2^ and host physiology^3^. This complex ecosystem has been extensively studied in the context of disease^4,5^, diet^6–8^ and age^9^ with the goal of determining how specific taxa and, more recently, the gene expression patterns of these taxa, impact human health. The relationship between the microbiome and the immune system has been of particular interest and specific bacteria have been shown to affect the function of both innate and adaptive immunity^10^. Further, an increasing number of inflammatory and autoimmune disorders have been associated with microbial dysbiosis^11^, though the precise mechanism for this relationship remains unclear.

Autoimmune diseases are multifactorial and chronic and the term covers nearly 100 distinct disorders^12^. Although there appears to be some genetic component, studies in disease-discordant twins have found that concordance rates are incomplete and therefore environmental factors, including the gut microbiome, likely contribute to disease pathogenesis^13,14^. Hundreds of studies have been carried out to better understand the connection between the microbiome and autoimmunity including studies specifically focused on inflammatory bowel disease (IBD), multiple sclerosis (MS), rheumatoid arthritis (RA), type 1 diabetes (T1D), and systemic lupus erythematosus (SLE). Despite the extensive study of the human gut microbiome in autoimmune disease, published results are inconsistent, which can be attributed to the differences in origin of samples (e.g. fecal or mucosal), sequencing platforms^1^, sample sizes, therapies administered, patients’ age^9^, geographical location^18^, and methods of data analysis. Thus, the question of whether there are common microbial features characterizing general autoimmunity still remains.

Therefore, to better understand the role of specific taxa in autoimmunity, we have reprocessed and reanalyzed 42 16S and metagenomic studies focused on the gut microbiome and autoimmunity. To do this, we have taken advantage of several machine learning approaches to provide an alternative to the traditional diversity analysis^19–21^. We specifically chose Random Forest (RF)^22^, eXtreme Gradient Boosting (XGBoost)^23^, Support Vector Machine^24^ with Recursive Feature Elimination^25^ (SVM RFE), and Ridge Regression^26^ algorithms since in addition to predicting a label they rank features according to how important the feature is for the label (disease) prediction. Random forest is a decision tree algorithm that has shown to be one of the most effective methods for classification of microbiome data, particularly 16S rRNA sequencing data.^19^ XGBoost, also a tree-based algorithm, has been recently shown to outperform other machine learning algorithms on a variety of biological datasets^27,28^. Further, we included ridge regression, another widely-used algorithm that differs from these tree-based models in that it is a logistic regression algorithm with L2 regularization that still enables us to compare its feature ranking to other algorithms. Finally, we used SVM RFE since it is a powerful feature selection method that has been used in numerous biomedical applications^29^. These methods give an advantage of learning functional relationships from the data without a need to define them beforehand. Moreover, many machine learning methods can handle sparse data with a large number of features, ranking them based on importance in their ability to distinguish between health and disease states^30^. These algorithms were used to identify microbial features predictive of general autoimmunity, as well as individual autoimmune diseases through the reanalysis of publicly available data on human gut microbiome in autoimmune diseases from the previous 10 years.

## MATERIALS AND METHODS

### Data acquisition

The NCBI PubMed database was searched for publications on April 1, 2020 related to the gut microbiome in autoimmune diseases from the last ten years based on the following criteria: 1) the study was performed on human fecal samples; 2) the subjects in the studies were older than 2 years old; 3) the samples were sequenced with either 16S rRNA sequencing or shotgun metagenomics or both; 4) the raw data in FASTQ format were publicly available; 5) the provided metadata allowed us to distinguish between disease and healthy control samples, as well as between subjects who were explicitly treated in the study and untreated samples. We identified a total of 42 studies, 30 with 16S rRNA sequencing data, 9 with shotgun metagenomics and 3 studies with both types of data available. In order to balance the number of the subjects with autoimmune disease with the number of healthy controls, we added 2 additional 16S rRNA studies, from which we selected only the healthy controls. Also, we included both 16S rRNA and shotgun metagenomics healthy samples from Human Microbiome Project 1 (HMP1) (**STable 1**).

### 16S rRNA data preprocessing

We employed QIIME2^31^ (v. 2018.11) to obtain the taxonomic abundances of the samples within each study, which were reprocessed independently and only the first time point was selected from each subject. Following data input, 454-based data underwent an error correcting step with *qiime dada2 denoise-pyro* command while the remaining samples were processed with either *qiime dada2 denoise-paired* or *qiime dada2 denoise-single* commands depending on whether the reads were paired or single (**STable 1**). During this process the bases with quality less than 20 were removed and the paired reads were merged. The resulted sequences abundance tables were rarefied to the depth of 5000. This depth was selected based on the alpha diversity curves of the studies, in which the plot reached the plateau. Further, we tried to account for 454-specific data since the sequencing depth of 454 samples was significantly lower than that of Illumina or Ion Torrent. As a result, the samples with sequencing depth less than 5000 were excluded from the further analysis (**SFig. 1**). In the next step we assigned the taxonomy to the sequences by training a Naïve Bayes classifier with *qiime feature-classifier fit-classifier-naive-bayes command* based on the Greengenes database^32^. Following taxonomy assignment, the taxonomic abundances tables were collapsed on both genus and species taxonomic levels. Further the resulting abundance tables from each study was merged together to create an “autoimmunity” data matrix or a disease-specific matrix.

### Shotgun metagenomics preprocessing

Sequencing reads were trimmed with Trimmomatic^33^ (v. 0.36) to have a quality of 20 or greater. KneadData^34^ was used to remove host sequences from reads, which were then supplied to the MetaPhlAn2^35^ to obtain relative taxonomic abundance, after which tables from individual studies were merged. One exception was the *Cekanavicute et al*. study, for which only preprocessed tables were available, which were processed in the same way.

### Predictive modeling

Caret package^36^ in R was used to build the predictive models which were built separately for each data type. For 16S rRNA we built 4 disease-specific models: autoimmune disease samples vs. healthy controls, IBD samples vs. healthy controls, MS samples vs. healthy controls and RA vs. healthy controls. We built those models on all samples that passed our inclusion criteria and on only adult (18 years and older) samples since children gut microbiome might exhibit higher interpersonal variation. In addition, we built predictive models comparing IBD and MS, IBD and RA, and MS and RA. For MS and RA models for which only adult samples were used. All models were trained on both genus and species taxonomic levels. Since we identified only 13 studies with publicly available shotgun metagenomics data, we computed only 2 metagenomics models: all autoimmune disease samples vs. healthy controls model and IBD vs. healthy controls model. Also, since there were significantly more healthy samples than disease samples, when considering the individual disease models, we randomly selected the same number of healthy controls samples to match the number of available disease samples. The data were split into training (90%) and test (10%) sets. The predictive models for each dataset were built with four models: Random Forest^22^, XGBoost^23^, Ridge Regression^26^ and SVM^24^ with radial kernel and RFE^25^ with a step of 2. Those models were selected due to their ability to rank the features based on the importance for the label prediction. To reduce the computing time before the training step the near-zero-variance features were identified and removed. In order to avoid overfitting, 5-fold-3-times cross-validation was employed to tune the models during the training step.

### Feature Selection

Each of the selected algorithms ranked features based on importance to their classification. Since the four algorithms employ different metrics for the feature ranking, first we sorted the features in the ascending order based on importance in each algorithm and then assigned the least important feature a value of 1, while the most important feature got the maximum score equal to the number of features in a given disease model. Then we selected the top 30 most important features for each disease model.

### Metabolomic analysis

We selected taxa that overlapped between at least one disease vs disease models, were identified on the genus level, and were present in the shotgun metagenomics dataset from The Inflammatory Bowel Disease Multiomics Database (IBDMDB)^37^. This method provided 12 different genera, 2 of which were filtered out due to study-based predictive power (**SFig. 7**). In the next step we correlated the abundance of the remaining 10 genera in the IBDMDB with the metabolomics table from IBDMDB by using pairwise Spearman correlation with Benjamini-Hochberg correction for multiple comparisons and selected metabolites based on correlations with an adjusted p-value cutoff of 0.05.

## RESULTS

### Autoimmunity-associated changes in microbial composition

We used a standardized meta-analysis approach to collect, reprocess and integrate available metagenomics data from case-control autoimmunity studies focusing on changes in the gut microbiome from human fecal samples. Using an expansive literature search we identified a total of 132 autoimmunity studies fulfilling our criteria (**SFig. 1**). Following filtering based on unique data, age (2 years or older), metadata and raw file availability and sequencing depth we were able to successfully download raw (FASTQ)16S rRNA and/or shotgun metagenomics data from 42 studies, 30 with 16S rRNA sequencing data^38–69^ and 9 studies with shotgun metagenomics data^37,70–77^, and 3 studies with both^78–81^ (**STable 1, SFig. 1**). These included studies on Inflammatory Bowel Disease (IBD, *N=14*), Multiple Sclerosis (MS, *N=7*), Rheumatoid Arthritis (RA, *N=5*), Juvenile Idiopathic Arthritis (JIA, *N=3*), Systemic Lupus Erythematosus (SLE, *N=3*), Type 1 Diabetes (T1D, *N=2*), Behcet’s Syndrome (BS, *N=2*),Ankylosing Spondylitis (AS, *N=2*), Antiphospholipid Syndrome (APS, *N=1*), Primary Sclerosing Cholangitis (PSC, *N=1*), Myasthenia Gravis (MG, N=1) and Reactive Arthritis (ReA, *N=1*) (**Fig. 1, SFig. 2**). Three additional studies with healthy subjects were included to balance the disease and non-diseased cohorts (**STable 1**).

**Figure 1:**
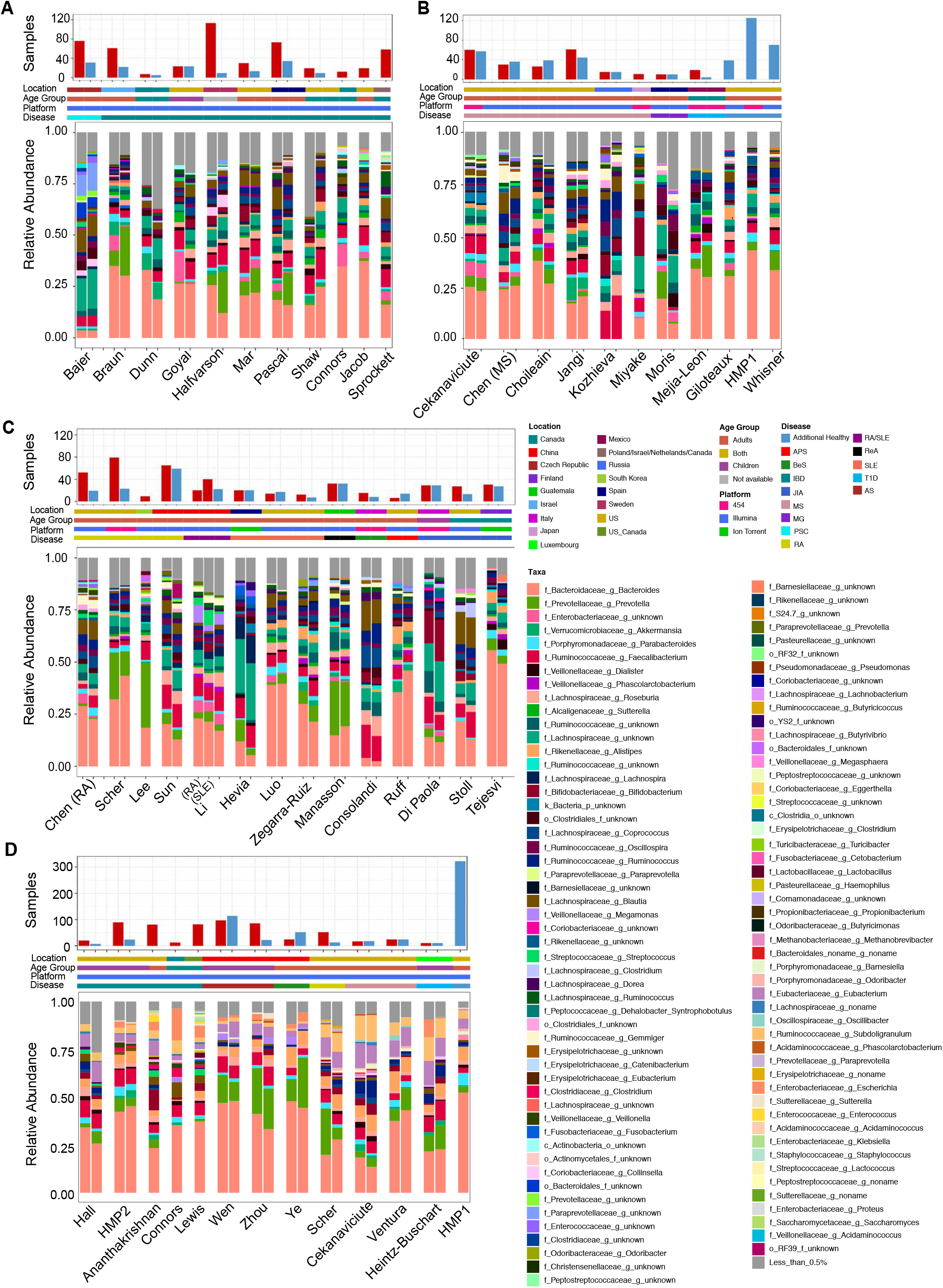
Autoimmunity Analysis Workflow. Thirty 16S rRNA sequencing datasets from studies focused on 12 different autoimmune diseases and two datasets representing healthy (non-autoimmune disease) cohorts were reprocessed using QIIME2^31^ with data2 denoising and rarified to 5000 sequence depth. The output species and genus level relative abundance matrices were used to create four machine learning models for (1) general autoimmunity; (2) inflammatory bowel disease (IBD); (3) multiple sclerosis (MS); and (4) rheumatoid arthritis (RA). Top ranked features from these models were identified and metabolic changes associated with these taxa of interest were assessed using the IBDMDB dataset^37^.

Initially, 16S rRNA data was reprocessed using a standard analysis pipeline, which included filtering and taxonomic assignment. Each study was reprocessed individually and final taxonomic abundance tables were then concatenated to a build a final autoimmunity matrix. Disease specific datasets were also created through combining reprocessed data tables for each individual disease type. Each table was then used to build predictive models of general autoimmunity as well as disease-specific models (**Fig. 1**) with the primary goal of identifying the most important features (taxa) involved in autoimmunity across and within disease types. Metagenomics data was also reprocessed using a separate analysis pipeline, providing taxonomic abundance tables (**SFig. 2**).

Following quality control (QC) and filtering, 30 16S rRNA^38–45,47–67,69,78,79,81^ and 12 metagenomics^37,70–79,81^ studies remained for downstream analysis (**Fig. 1, SFig. 2**). Notably, 10 out of the 30 16S rRNA and 5 of the 12 metagenomics studies used investigated the role of the human gut microbiome in IBD, due in part to its relatively high prevalence in 1.3% of US adults^82^. However, we were also able to acquire data from studies of more rare autoimmune diseases including Behçet’s Syndrome, which results from inflammation of the blood vessels^43^, Myasthenia Gravis, a neuromuscular disorder, and Reactive Arthritis. A portion of these studies contained significantly more disease samples than the healthy samples, with Halfvarson *et al*. having 10 times more samples from individuals with autoimmune disease than from healthy controls, and with 6 other studies^50,53,64,71,72,79^ containing only disease samples (**Fig. 2**). For this reason we included healthy samples from three additional studies which investigated non-autoimmune diseases^46,68,80^, which after QC and preprocessing resulted in additional 232 16S and 322 shotgun metagenomics samples.

**Figure 2.**
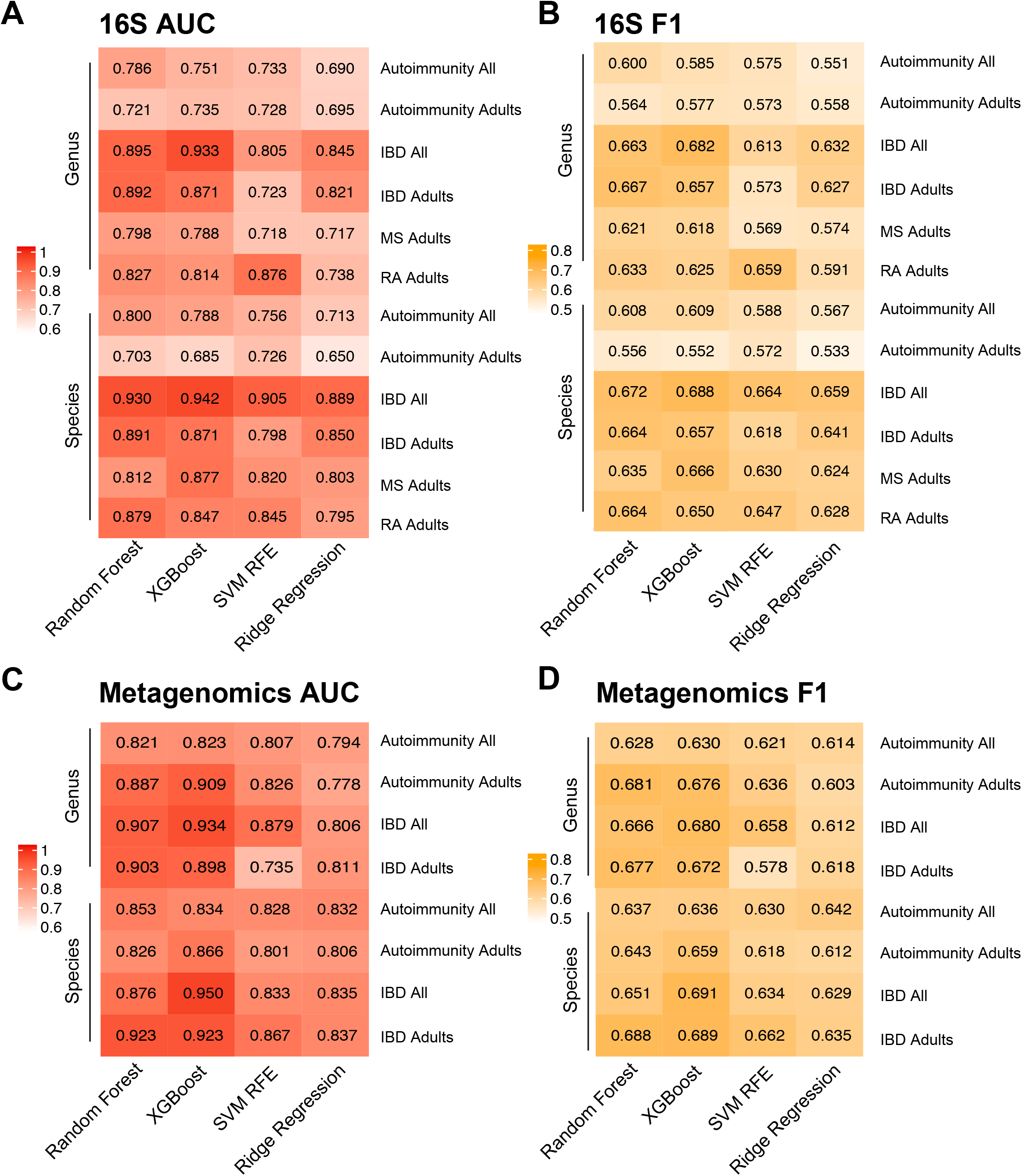
Study overview. Overview of (a) 16S rRNA sequencing and (b) shotgun metagenomics studies included in our analysis. Includes the number of healthy (blue bars) and disease (red bars) samples in each, geographic location, age group and disease studied. Also includes the average relative abundance of taxa at the genus level for the healthy and diseased subjects following re-processing. Inflammatory bowel disease (IBD), multiple sclerosis (MS), rheumatoid arthritis (RA), juvenile idiopathic arthritis (JIA), systemic lupus erythematosus (SLE), type 1 diabetes (T1D), primary sclerosing cholangitis (PSC), Behcet’s syndrome (BeS), ankylosing spondylitis (AS), antiphospholipid syndrome (APS), myasthenia gravis (MG) and reactive arthritis (ReA).

While combining these diverse datasets there were several study-specific characteristics known to impact microbial identification that we paid specific attention to, such as geography, age, sequencing platform and 16S rRNA primers. A majority of the studies were based on populations from North America, Europe and Asia, however Manasson *et al*. investigated the gut microbiome of spondyloarthritis in Guatemalan patients^56^, Mejia-Leon *et al*. looked at Type 1 Diabetes in Mexico^58^, while Sprockett *et al*. had participants from four different countries, Poland, Israel, Netherlands and Canada^64^. Further, there was a large range in age across studies, with participants being from 2 to 76 years old. Studies focusing on newborn children (less than 2 years of age) were not included since it has been well established that the microbial diversity in the first few years of life is significantly lower when compared with adults^83^. We also controlled for age by building separate models for adults only (18 years or older) in addition to models including all participants in datasets where children were included (general autoimmunity and IBD). DNA was sequenced with one of three sequencing platforms, 454 pyrosequencing, Ion Torrent, or Illumina instruments with both paired and single reads techniques. Description of the characteristics for each study can be found in **Supplemental Table 1** and **Figure 2**. To assess potential batch effects, we employed a Principal Coordinate Analysis (PCoA)^84^ based on the Bray-Curtis distance^85^ and investigated non-disease based differences. No significant differences were observed based on autoimmune disease diagnosis, however, there were a subset of non-disease characteristics that were identified as significant based on study, subject characteristics, or sequence methods (**SFig. 3, SFig. 4**). To combat this, we completed study based analysis to identify study-specific vs. disease-specific features as part of our downstream analysis (**SFig. 7**).

We first examined the taxonomic composition on the genus level of the healthy and diseased samples in each study to verify expected changes based on previously published results. We were able to recapitulate major findings from all studies. For example, we identified disease-specific alterations in multiple studies in *Akkermansia*^86,51^, *Bacteroides*^45,49,74^, *Blautia*^55,56^, *Clostridiaceae*^44^, *Faecalibacterium*^65,41^, *Lachnospira*^48,65,57^, *Parabacteroides*^78^, *Prevotella*^56–58,73,74^, *Ruminococcacaea*^48,57,86,56,45^ and *Streptococcus*^41^ (**Fig. 2**). Interestingly, these previously published results, and our reanalyzed results, varied in the directionality of the change for many of these taxa, with disease specific overabundance occurring in a subset of studies and a reduction in other. These inconsistencies further highlight the need for standardized reanalysis and integration of these valuable datasets to better understand the potential impact of microbial changes in autoimmune disease.

The taxonomic composition of healthy individuals showed clear differences, which can be attributed to several factors. First, it is well established that microbial composition differs by age and geography^18^. Secondly, it is not guaranteed that the “healthy” recruits included in these studies did not suffer from another pathology impacting the gut microbiome. In most studies, researchers only ensured that healthy controls had not been diagnosed with an autoimmune disease of interest and had not taken antibiotics at least during the sample collection. Thirdly, as these studies were sequenced on different platforms and with differing 16S rRNA hypervariable regions during PCR amplification, we expect a level of variability in the identified taxa even across controls^87^.

### Predictive Modeling of Autoimmunity

In order to identify which taxa are most important for distinguishing between healthy controls and subjects with autoimmune disease we built four independent machine learning disease models on 16S rRNA data: (1) IBD specific; (2) MS specific; (3) RA specific; and (4) general autoimmunity; which included samples from all the autoimmune diseases available (**Fig. 1**). Genus level taxonomic abundances were used for the final predictive modeling analyses. Four independent algorithms were used to capitalize on the strengths and limitations of each: Random Forest (RF)^22^, eXtreme Gradient Boosting (XGBoost)^23^, Support Vector Machine^24^ with Recursive Feature Elimination^25^ (SVM RFE), and Ridge Regression^26^. For both the general autoimmunity and IBD model, an ‘Adult only’ model was also created, removing all participants younger than 18 years old, to control for known age-specific differences in microbial composition. MS and RA models included only adults. Application of four independent algorithms capable of feature ranking to the same data provided an advantage in robustly identifying the most important features predictive of autoimmunity by multiple models, providing an additional level of confidence. Models were run at both the genus and species level.

Model performance was evaluated using both Area Under the receiver operating characteristics Curve (AUC) and macro F1 score, which reports on the reciprocation between the specificity and sensitivity. Notably, we incorporated near-zero-variance feature removal to reduce both computational load and to consider only features with reasonable variation between the samples, as those with little variation likely would not impact disease state. Among the four algorithms for the autoimmunity model, the best performance was achieved by Random Forest with an AUC of 0.8 using the species level data. The superior performance by this algorithm was not unexpected, as Random Forest has been previously shown to perform well on microbial data^19^. Random Forest was also the best predictor for the species-level RA model, with an AUC of 0.879. XGBoost produced the best AUC for the IBD and MS disease prediction of 0.942 and 0.877 respectively (**Fig. 3**). In general, model performance was similar at the species and genus level, with slightly higher AUCs occurring in the species models. In addition, we applied the same predictive modeling strategy to shotgun metagenomics data. Due to data availability, we built only general autoimmunity and IBD models, with highest AUCs for ‘Adult only’ models reaching 0.877 for the general autoimmunity model and 0.923 for the IBD model.

**Figure 3.**
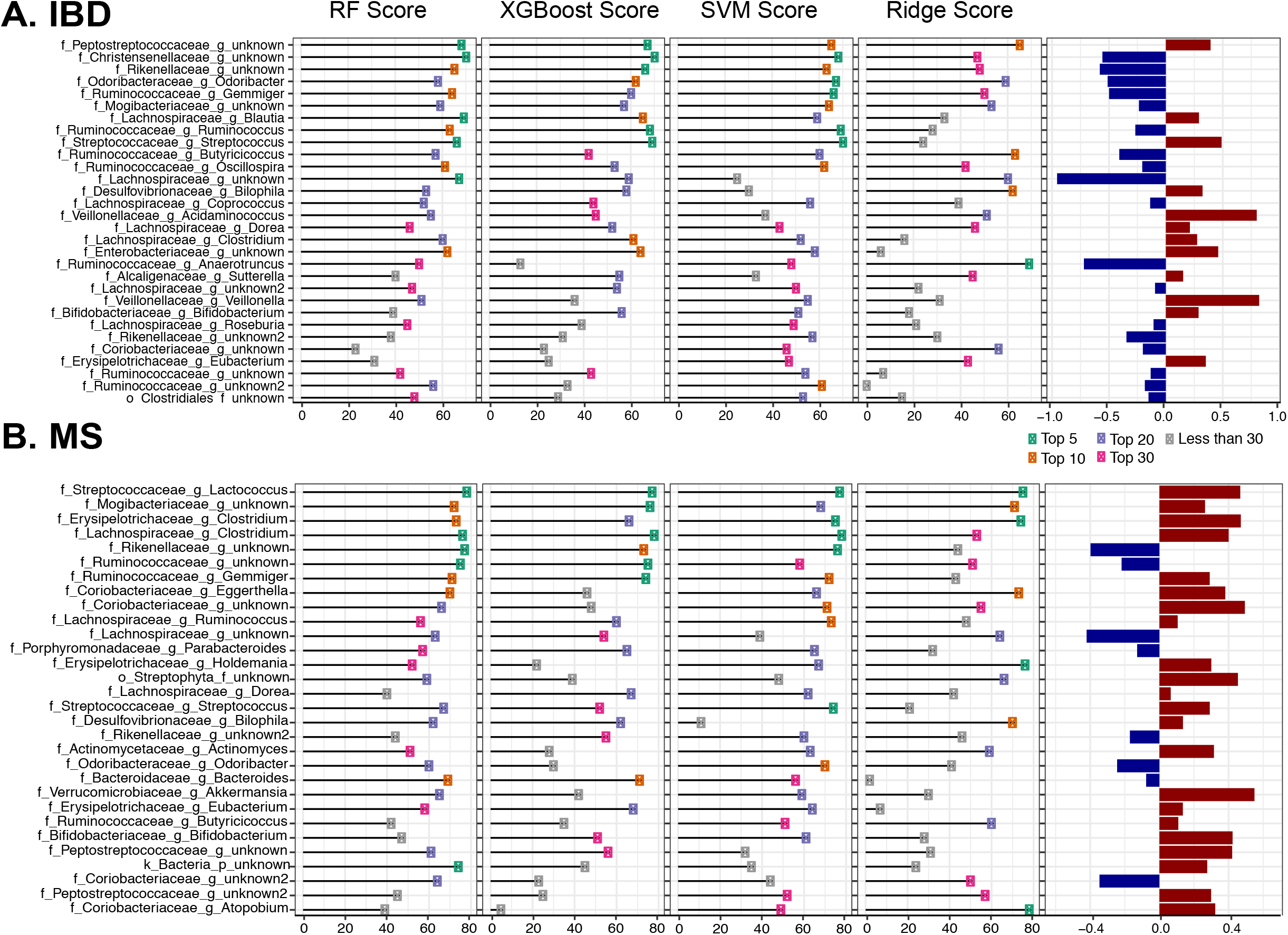
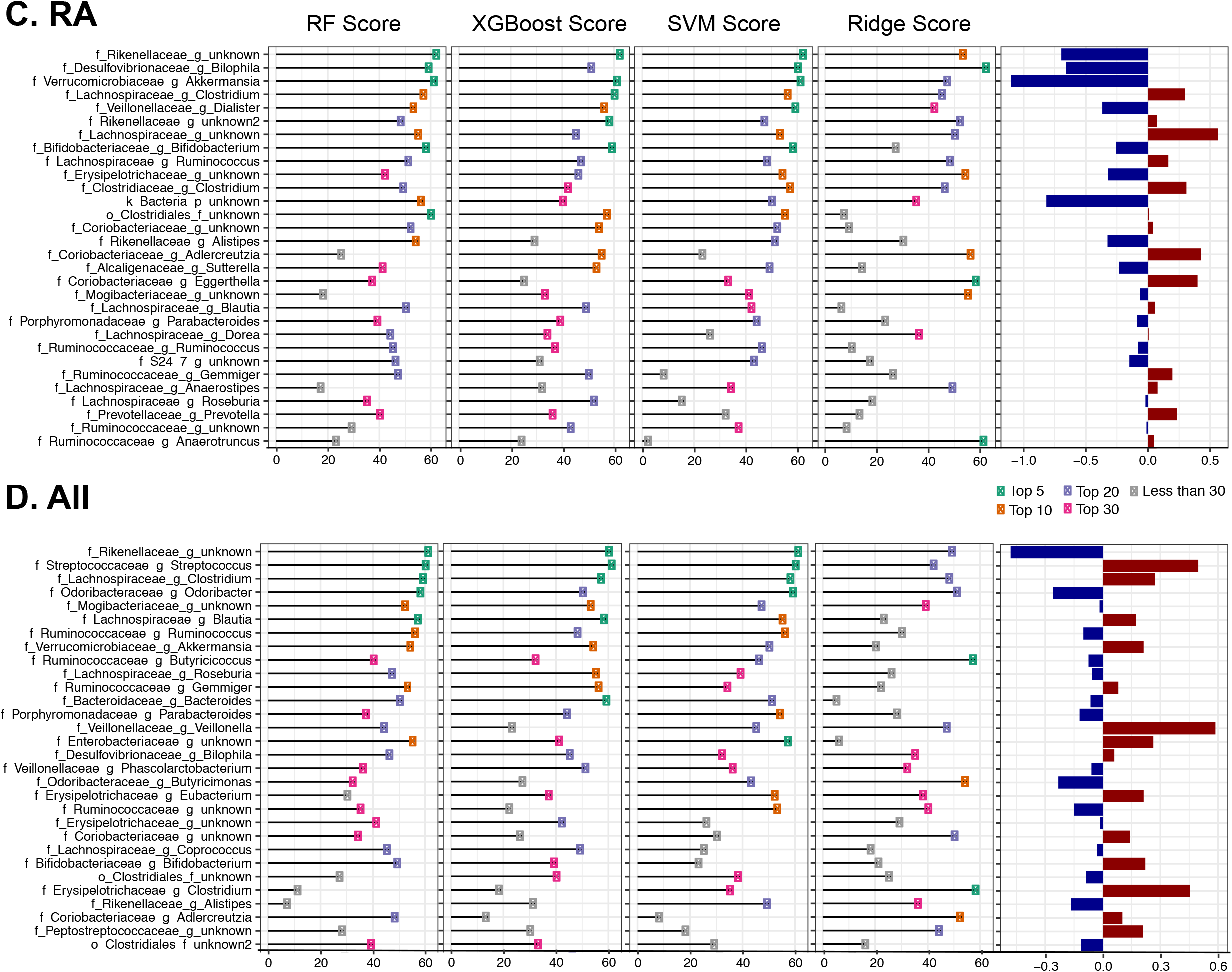
Predictive Modeling of Autoimmune Disease. Area under the curve (AUC) (a,c) and F1-scores (b,d) for models predicting any autoimmune disease, inflammatory bowel disease (IBD), multiple sclerosis (MS) and rheumatoid arthritis (RA) for four machine learning models, random forest, XGBoost, support vector machine with recursive feature elimination (SVM RFE) and ridge regression and at the genus and species level. Adult-only (18 years and older) and All (adult + children) models are included for general autoimmunity and IBD studies.

Overall, the most stable AUC across the three algorithms was reached on the IBD data set, likely due to the considerably higher number of IBD samples compared with other autoimmune diseases. Notably, we were able to predict autoimmunity based on only microbial composition of the samples, which suggests that there exists a common gut microbiome signature present that may be relevant to all autoimmune diseases. In order to determine whether our AUCs could be predicted by chance, we assigned the labels to the samples at random, and computed our models again. The models trained with the random label assignment produced the AUCs of ~0.5 (**SFig. 5)**, which is indicative of a true difference between the healthy controls and autoimmune disease subjects based on the gut microbial composition.

### Most Predictive Model Features

Since all four of our models employed feature ranking we were able to identify which features were most important for predicting the three distinct autoimmune diseases as well as general autoimmunity. From this, we identified features that were ranked similarly by all four algorithms. The top 30 features were selected based on a combined feature score of ranked taxa across all 4 models for each disease (**Fig. 4, SFig. 6**). In order to account for potential batch affects occurring due to study population differences (**SFig. 3**), we created “mock” models to predict the study a sample came from, regardless of disease status. This allowed us to identify taxa that were able to specifically identify a study population rather than the disease. These models identified *Coriobacteriaceae, Bacteroidales, Rikenellaceae, Stretococcaceae Streptococcus, Lachnospiraceae Blautia, Lachnospiraceae Dorea, Alcaligenaceae Sutterella* and *Enterobacteriaceae* as able to predict study regardless of disease or healthy status in at least one study (**SFig. 7**). This allowed us to identify taxa that are likely tied to the study population, sequencing platform or experimental method, rather than disease status.

**Figure 4.**
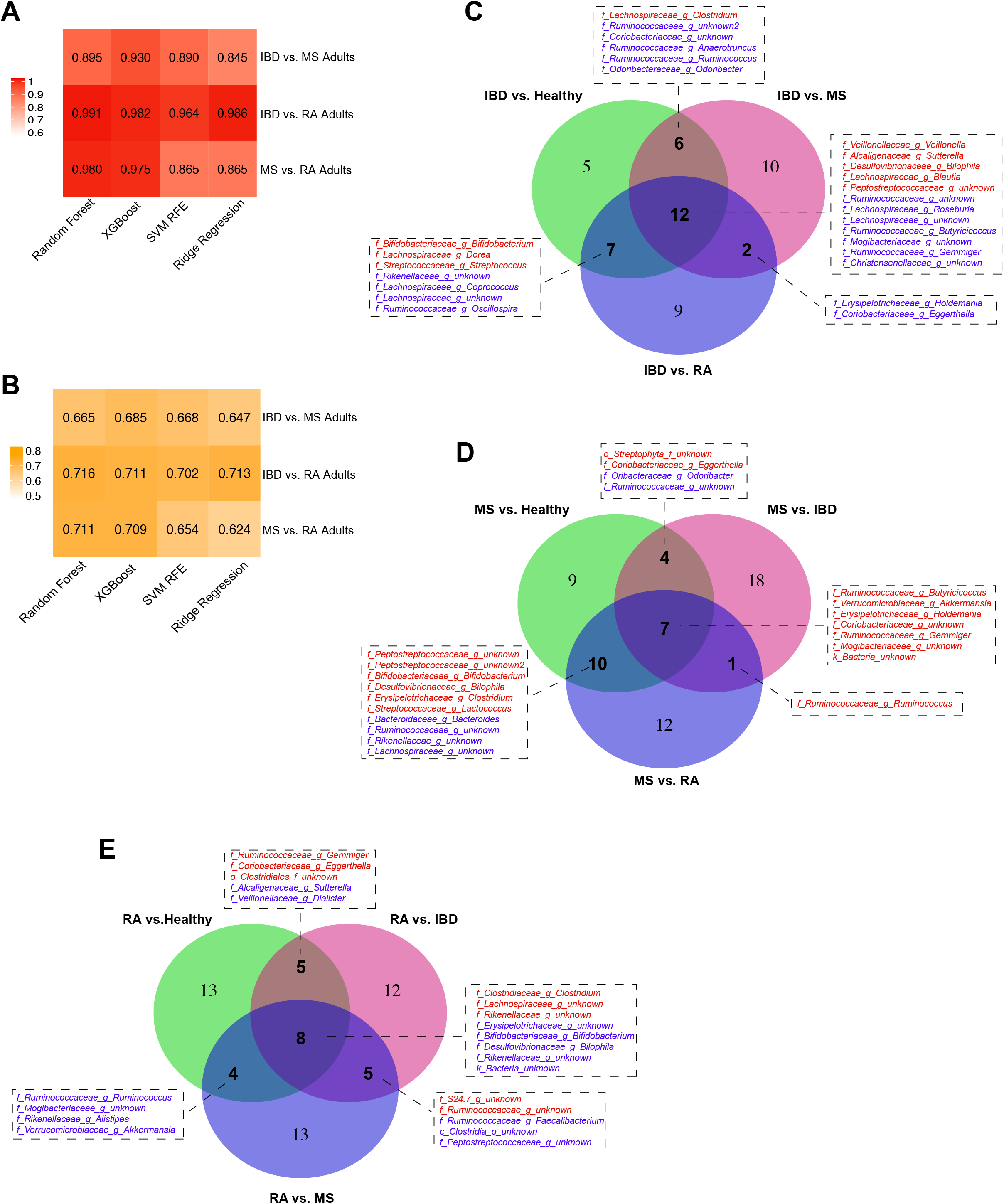
Taxa Predictive of Disease. Top 30 taxa across four predictive models (adults only), random forest (RF), XGBoost, and support vector machines (SVM), ridge regression for (a) inflammatory bowel disease (b) multiple sclerosis, (c) rheumatoid arthritis and (d) general autoimmunity. Features ranked by mean rank across the four models and color indicates the rank of each taxa in each model. Log fold change of disease vs. healthy for each taxon.

The most predictive features identified by our IBD model were reduced levels of *Christensenellaceae, Odoribacter* and *Gemmiger* and increased abundance of *Peptostreptococcaceae* and *Lachnospiraceae Blautia* (**Fig. 4a**). MS predictive features included increases in *Lactococcus, Mogibacteriaceae, Erysipelotrichaceae Clostridium* and *Lachnospiraceae Clostridium* and reduced levels *Ruminococcaceae* (**Fig. 4b**). Further, the RA model identified reduced abundance of *Desulfovibrionaceae Bilophila, Akkermansia* and *Veillonellaceae Dialister* and increased levels of *Lachnospiraceae Clostridium* as most predictive of disease state (**Fig. 4c**). Lastly, for our comprehensive autoimmunity analysis, we identified *Odoribacter* and *Mogibacteriaceae* as the most important features with reduced abundance in autoimmune disease samples compared with healthy controls and *Streptococcus* and *Clostridium having* increased expression in diseased participants (**Fig. 4d**). Although *Rikenellaceae* was repeatedly identified by all models, our study-specific models also identified this genus as being study specific and therefore we did not consider it in our downstream biological interpretation (**SFig. 7**)

By also comparing our three disease types (IBD, MS and RA) to each other we were able to further refine our disease specific predictive taxa from our heterogeneous dataset. To do this, we again used predictive modeling (Random Forest) to compare each disease to each other, identifying a new set of predictive taxa, and overlapped these with those identified in the original model created based on healthy controls. The model performance (AUC, F1 score) and overlap of the thirty most predictive taxa from each model is shown in **Figure 5**. This analysis provided us with a list of taxa able to distinguish each disease not only from healthy controls, but from other autoimmune diseases. In IBD, 12 features were identified in all three comparisons, including increased *Peptostreptococcaceae*, and decreased levels of *Mogibacteriaceae* and *Gemmiger* (**Fig. 5c**). Increased *Butyricicoccus, Akkermansia, Holdemania* and *Odoribacter* were four of the seven taxa consistently predicted in our MS models (**Fig. 5d**) and increased *Clostridiaceae Clostridium*, and reduced *Erysipelotrichaceae*, and *Desulfovibrionaceae* were three of the eight identified in all RA models (**Fig. 5e**).

**Figure 5.**
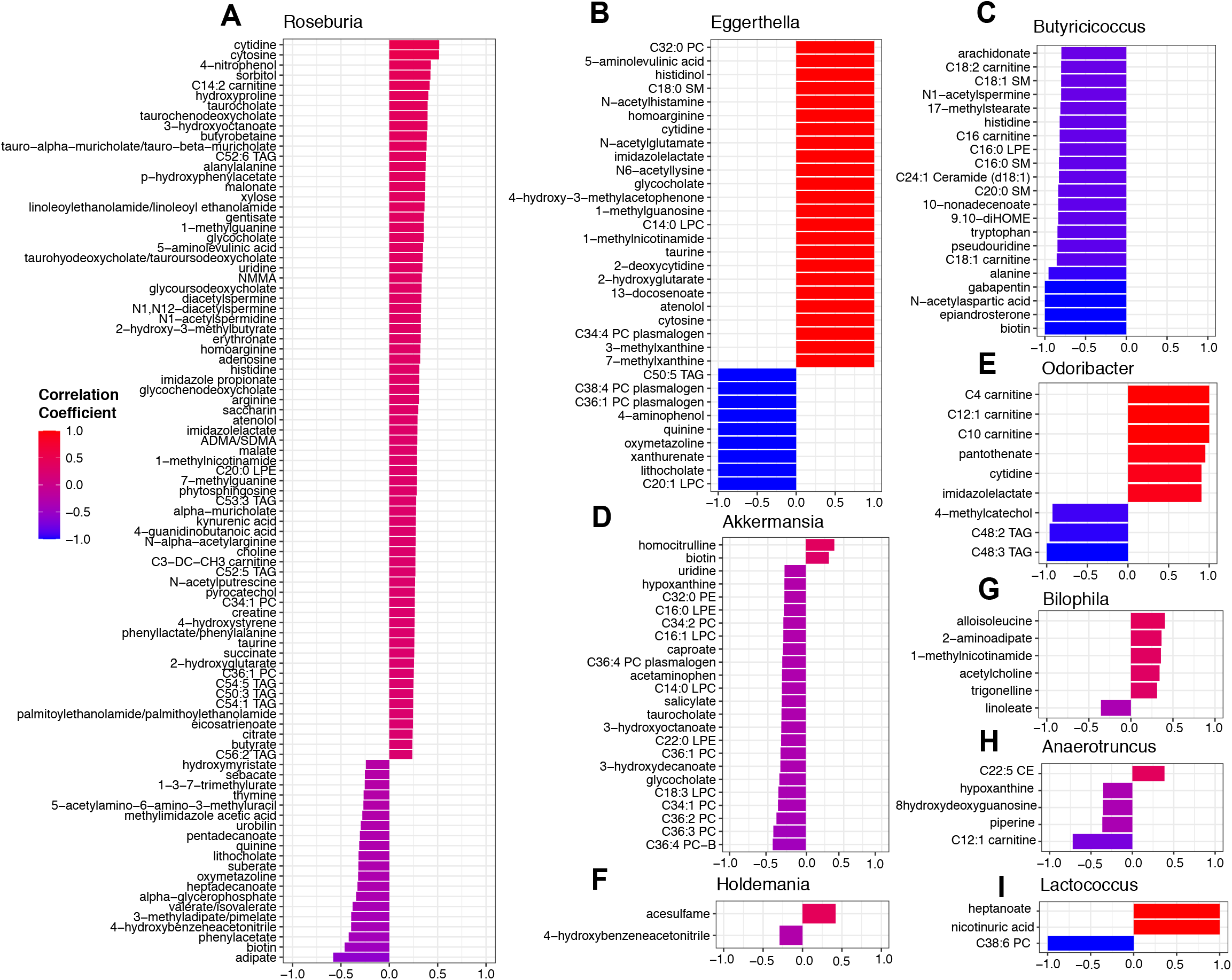
Disease vs. Disease Comparison Models. Model (a) AUCs and (b) F1 scores when predicting diseased samples when compared against other disease. Taxa consistently identified in multiple comparison models for (c) inflammatory bowel disease (IBD), (d) multiple sclerosis (MS), and (e) rheumatoid arthritis (RA).

To validate these findings, we also applied the same machine learning approach to shotgun metagenomics data from 13 studies^37,70–81^ (**SFig. 2**). Six of the top 15 features most predictive features overlapped in both the 16S autoimmunity (**Fig. 4a**) and metagenomics autoimmunity adult models (**SFig. 6e**), including *Clostridium, Odoribacter and Parabacteroides*.Similarly, both 16S (**Fig 4b**). and metagenomics (**SFig. 6f**) IBD models had 3 overlapping top features including *Odoribacter* and *Ruminococcus*.

### Correlations between highly ranked taxa and metabolism in IBD

To better understand the potential downstream effects of altered abundance levels of these taxa, we used the Inflammatory Bowel Disease Multiomics Database (IBDMDB) metabolomic dataset to identify metabolites which are significantly correlated with our taxa of interest. For this purpose, we chose features that overlapped in at least two of the three disease vs disease models that identified on the genus level (25 taxa total, **Fig. 5c-e**) and which were present in the IBDMDB shotgun metagenomics dataset. This resulted in a total of 12 genera in common between our dataset and IBDMDB cohort (**Fig. 6, SFig. 8**). Two of these taxa (*Lachnospiraceae Dorea, Alcaligenaceae Sutterella*) were excluded from this analysis as they were also flagged as being consistently able to predict study regardless of disease or healthy status of the samples (**SFig. 7**).

**Figure 6.**
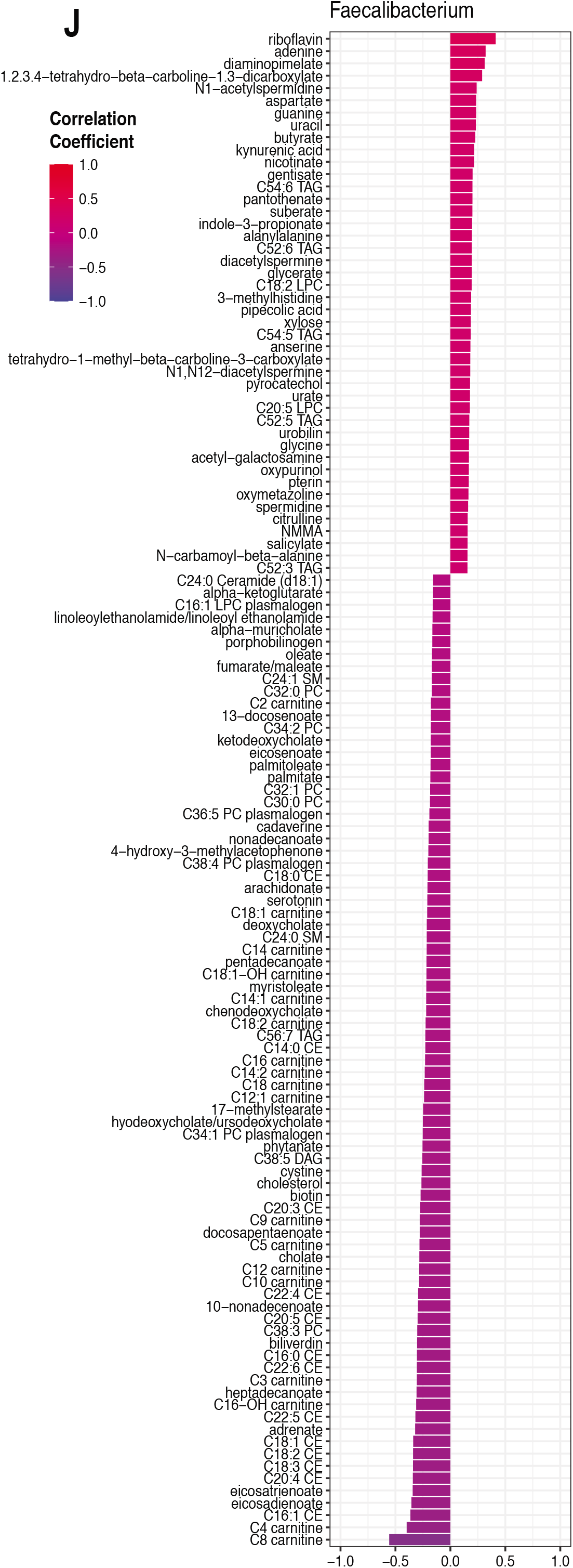
Metabolites significantly correlated with disease-predictive taxa. Spearman correlation coefficient scores plotted and shaded by adjusted p-value for 10 taxa found to be predictive of IBD, MS and RA based on the multiple disease model comparisons.

Investigating correlations between the abundance of the remaining 10 genera with metabolites within the IBDMDB, we identified 250 metabolites that significantly correlated with at least one taxon at an adjusted p-value < 0.05. Two of the 10 genera assessed, *Roseburia* and *Butyricicoccus*, were found to be reduced in IBD (**Fig. 5c**). *Roseburia* had the highest correlations occurring in a number of bile acids (e.g. taurocholate, taurochenodeoxycholate, glycocholate), in addition to several triacylglycerols (TAGs) and fatty acids (**Fig. 6a**). Specifically, the short chain fatty acid (SCFA) butyrate was found to be positively associated with *Roseburia* abundance. Bacteria fermentation of carbohydrates in the gut are known to produce SCFAs and butyrate, in particular, is known for playing a critical role in host metabolism^88^ and intestinal inflammation through regulatory T cell (Treg) and T helper cell 17 (Th17) response^89,90^. Consistent with its known anti-inflammatory role in the intestine^91^, butyrate was associated with taxa having low abundance in IBD. The second IBD predictive taxon, *Butyricicoccus*, was associated with reduced levels in three long chain acylcarnitines, three sphingomyelins (C16:0, C20:0 and C18:1), arachidonate and C24:1 ceramide (**Fig 6c**).

Increased abundance of *Akkermansia, Lactococcus* and *Holdemania* were found to be predictive of MS (**Fig. 5d**). *Akkermansia* showed negative associations with the bile acid components taurocholate, bile acid glycocholate and fatty acid anions 3-hydroxyoctanoate and caproate (**Fig. 6d**). The identification of bile acids associated with a number of our taxa is consistent with several studies showing an integral role of the gut bile acid pool as a modulator of host immune response and inflammation^92,93^. *Holdemania* had increased levels of the artificial sweetener, acesulfame, and the benzyl cyanide, hydroxybenzene acetonitrile (**Fig. 6f**) while *Lactococcus* was positively associated with nicotinuric acid, a niacin metabolite (**Fig. 6i**). Interestingly, niacin has also been recently shown to attenuate gut inflammation in rodent models^94^.

*Odoribacter* was reduced in both MS and IBD models, and showed positive associations with several acylcarnitines, including C10 carnitine and C12:1 carnitine, which were both found to be significantly increased in subjects with dysbiotic Crohn’s disease^95^. As *Odoribacter* was found to be reduced in our IBD population, this is a somewhat contradictory finding, but may be due to the variability and unknown levels of dysbiosis in our population. Pantothenate (vitamin B5) was also found to be associated with this taxa, consistent with a recent study showing a depletion of pantothenate in the gut of IBD subjects^95^ as we would expect to see reduced levels of vitamin B5 with reduced *Odoribacter* levels.

Lastly, *Bilophila* and *Eggerthella*, were both found to be increased in RA (**Fig. 5e**). *Eggerthella* was found to be significantly associated with histamine pathway metabolite N-acetylhistamine and the bile acid lithocholate (**Fig 6b**). *Bilophila* was found to be positively associated with the branched chain amino acid alloisoleucine and the lysine metabolite, 2-aminoadipate.

## DISCUSSION

In this analysis, we used data from 42 studies investigating the role of the human gut microbiome in autoimmune disease, assessing both general autoimmunity and specific diseases. Since it is not always possible to find consistent differences using traditional metaanalysis methods, we applied classification algorithms to predict whether a sample comes from a healthy control or an autoimmune disease sample across multiple studies. We specifically used random forest, XGBoost, ridge regression and SVM RFE as these algorithms are capable not only predicting the disease status of the samples, but also ranking the features based on how important they are for the prediction.

Random forest has a long history in microbiome studies and has proven to be a robust algorithm that performs well on sparse unbalanced datasets. XGBoost is quickly gaining popularity and uses both types of regularization: L1 and L2 which prevents model from overfitting and it has in-built capability of handling sparse data and missing values. While random forest employs a bagging strategy, where each tree is provided with a full set of features and a sample of the data with replacement, XGBoost uses a boosting strategy, which is based on sequential training of shallow trees where each tree tries to correct the errors by the previous trees. Interestingly, both algorithms showed similar performance on our datasets. Further, we applied SVM RFE with a radial kernel, an algorithm that defines a non-linear hyperplane that maximizes the boundary between the two classes. In addition, SVM RFE utilizes recursive feature elimination, which is a wrapper algorithm that starts by training the model with all the features where the least important feature is eliminated, and it repeats this process until the best performance is reached. Due to the need to train the model repeatedly, SVM RFE requires significantly more training time than a regular SVM. For comparison we also used ridge regression, which is logistic regression with L2 regularization. Despite being a simpler model capturing only linear relations, it produced comparable results to the other three algorithms. A recent paper by Topcuoglu et al. evaluating the application of 7 diverse machine learning models to microbiome data found similar results to ours, with tree-based models performing best but with logistic regression with L2 regularization closely following^96^.

Connections between the gut microbiome and general autoimmunity have been made by studies investigating the role of human leukocyte antigen (HLA) gene polymorphisms in autoimmunity risk in a number of diseases including type 1 diabetes^97^, spondyloarthritis^98^, Behcet’s disease^99^, and Celiac disease^100^ explained through the impact of HLA on the amino acid sequence in class II major histocompatibility complex (MHC). It has been hypothesized that these polymorphisms may be involved in immune response in the gut and could be a link between autoimmune disease and the microbiome composition^101^. Interestingly, one of the top features identified by our IBD model, *Peptostreptococcaceae* (**Fig. 4**), was also identified as being associated with HLA risk alleles in a T1D risk study. This taxa was found to be significantly associated with a lower HLA genetic risk of autoimmunity, identifying it as a potential environmental trigger for autoimmune disease and warranting further study in IBD genetic risk based on our results^101^. We also identified a number of additional genera that were consistently predictive of disease. For example, reduced levels of *Lachnospiraceae Clostridium* and *Mogibacteriaceae* were identified as a top feature in all four of our disease models and serve as possible factors further connecting the gut microbiome and autoimmunity (**Fig. 4**).

In addition to identifying taxa predictive of general autoimmunity we were able to identify a number of novel taxa specific to IBD, MS and RA. Although several of these taxa have been previously associated with these diseases, conflicting and inconsistent results have been common. To try to circumvent these limitations, we have reanalyzed a large number of available gut microbiome studies to provide a broad perspective on the connection between the microbiome and specific disease. Our analysis has recapitulated several recent articles connecting the microbiome with autoimmunity and has also identified a number of novel taxa that may be related to these pathologies. For example, we found a depletion in *Roseburia, Ruminococcaceae* in IBD compared with controls, consistent with other studies of IBD^4^ and identified *Akkermansia* as a consistently predictive taxa for MS, an organism which has been shown to interact with spore-forming bacteria to worsen the impact of MS-associated microbiota^102^.

Further, many of the taxa we identified as being predictive of autoimmune disease were correlated with metabolites that have been potentially involved with autoimmunity and inflammation. Recent publications have identified a number of bile acids^92,93^, triacylglycerols^103^, vitamin B^94,95^, and acylcarnitine^95^ metabolites involved immune response and the microbiome, many of which we also found to be significantly associated with our most predictive taxa. Histamine, along with taurine and spermine which were also highlighted by our analysis, have been found to help shape the host-microbiome relationship through the regulation of the NLRP6 inflammasome signaling^104^. Further, we identified an association between IBD predictive taxa, *Roseburia*, with the SCFA butyrate, which among other SCFAs has been shown to inhibit histone deacetylases (HDACs) and inhibit immune response through Treg regulation and as ligands for G-protein coupled receptors with downstream anti-inflammatory effects^90,105,106^. The association identified between metabolites and taxa could be either due to the impact of that metabolite on the growth of the taxa, the metabolite being a produced by said taxa, or the metabolite negatively associating growth of an inhibitory species, and thus must be followed up by a more targeted approach to understand the precise biological mechanism.

Duvallet et al., completed a similar meta-analysis study in 2017 looking across 10 disease types (arthritis, autism spectrum disorder, Crohn’s disease, *Clostridium difficile* infection, liver cirrhosis, colorectal cancer, enteric diarrheal disease, HIV infection, liver diseases, minimal hepatic encephalopathy, non-alcoholic steatohepatitis, obesity, Parkinson’s disease, psoriatic arthritis, rheumatoid arthritis, type I diabetes and ulcerative colitis) to identify disease-specific and shared taxa^4^. They too, identified a number of genera associated with more than one disease, including *Lachnospiraceae and Ruminococcaceae* families and several members of the *Lactobacillales* order and showed the strengths of cross disease comparison using publicly available data. Studies delving into specific disease subcategories, such as this study focused on autoimmune disease, build upon their original study. Further, our reanalysis focused more acutely on investigation of inter-study batch effects and methods of reducing the impact of these on downstream analysis.

We understand there are several limitations of this study. Firstly, the sample size is relatively small for machine learning reducing model reliability. As additional data is generated on larger cohorts from different ages and different cultural backgrounds we can continue to develop and run similar models to further elucidate how gut microbiome promotes autoimmune diseases. Additionally, the differences in sequencing platform, geography and subject characteristics provide confounders that are difficult to remove from the dataset *post hoc*. Cautious evaluation of taxa identified by our methods in addition to the use of control models testing the ability to predict by study rather than disease were used to combat this issue, however we are aware that these confounders remain. Future analysis further evaluating how each of these study design techniques and participant make-up effects the results of a microbiome study would be of great benefit to the community.

---

**Supplementary Figure 1:** Study collection and filtering.

**Supplementary Figure 2.** Metagenomic analysis workflow.

**Supplementary Figure 3:** PCoA diagrams and statistical differences across 16S datasets showing sample similarity by (a) health status, (b)16S rRNA region sequenced, (c) sequence platform (d) disease type, (e) age group, (f) exact age, (g) country, (h) study, (i) antibiotics consumption, and (j) DNA extraction kit.

**Supplementary Figure 4.** PCoA diagrams and statistical differences across metagenomics datasets showing sample similarity by (a) health status, (b) age group, (c) disease type, (d) exact age, (e) country, (f) antibiotics consumption, (g) study, and (h) DNA extraction kit.

**Supplementary Figure 5.** AUCs for models trained with random label assignment for (a) 16S and (b) metagenomics studies.

**Supplementary Figure 6.** Top 30 taxa across four predictive models for 16S studies describing (a) general autoimmunity (adults + children), (b) inflammatory bowel disease (adults + children), and metagenomics studies investigating (c) general autoimmunity (adults + children), (d) inflammatory bowel disease (adults + children), (e) general autoimmunity (adults only), and (f) inflammatory bowel disease (adults only). Features ranked by mean rank across the four models and color indicates the rank of each taxa in each model. Log fold change of disease vs. healthy for each identified taxon is also shown.

**Supplementary Figure 7.** Models and features predictive of study. AUCs of random forest model prediction of study and taxa features most predictive of study in those models in 16S (a,c) and metagenomics (b,d) studies.

**Supplementary Figure 8.** Significant correlations between metagenomic abundance of 10 selected genera and metabolites in IBDMDB dataset.

## TABLES

**Supplementary Table 1:** Study details including, sample number, 16S rRNA region, and sequencing platform.

