## Supplementary figures and images for "Predictive Metagenomic Analysis of Autoimmune Disease Identifies Robust Autoimmunity and Disease Specific Microbial Signatures"

### Supplemental Figure 1

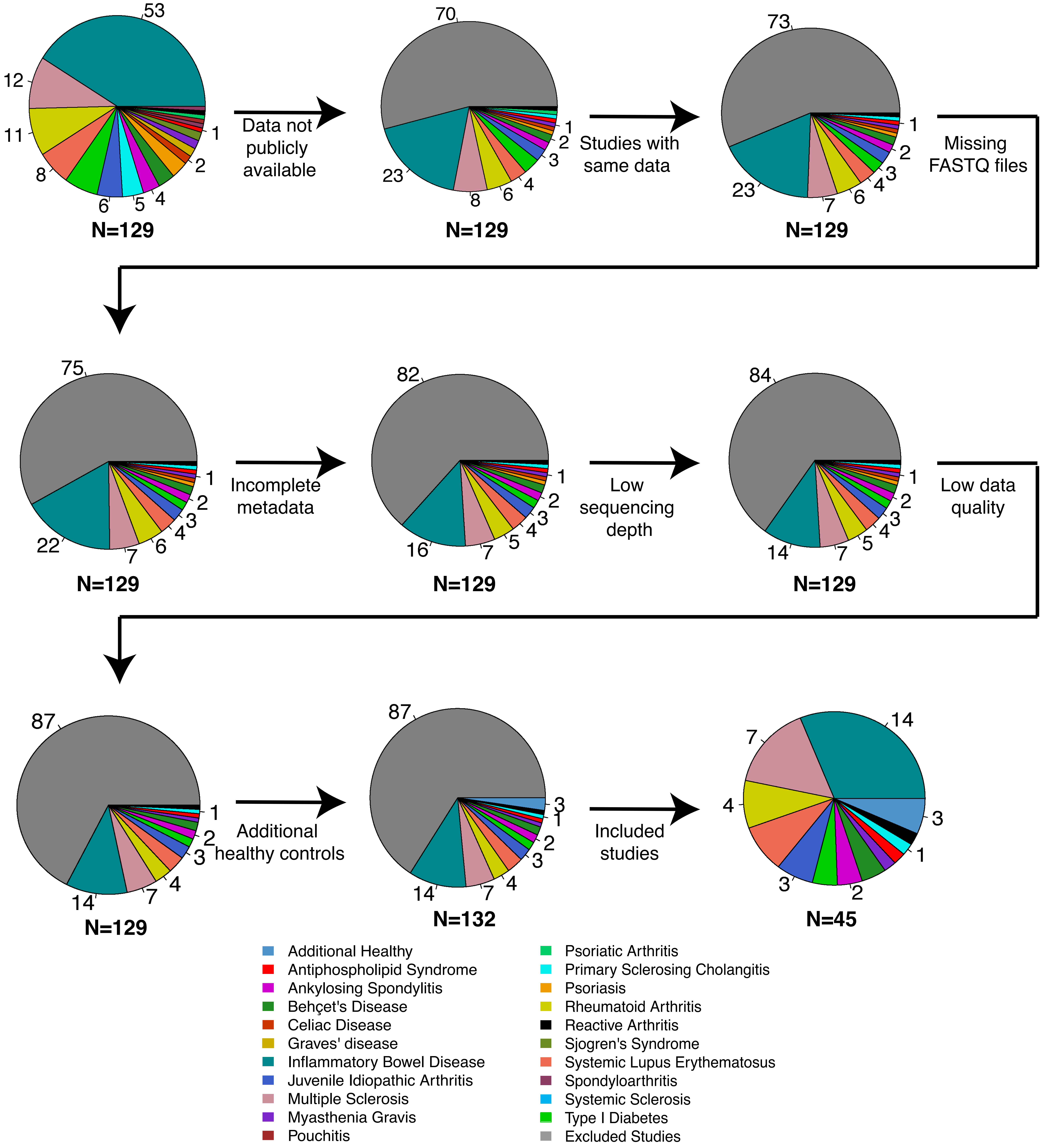

### Supplemental Figure 2

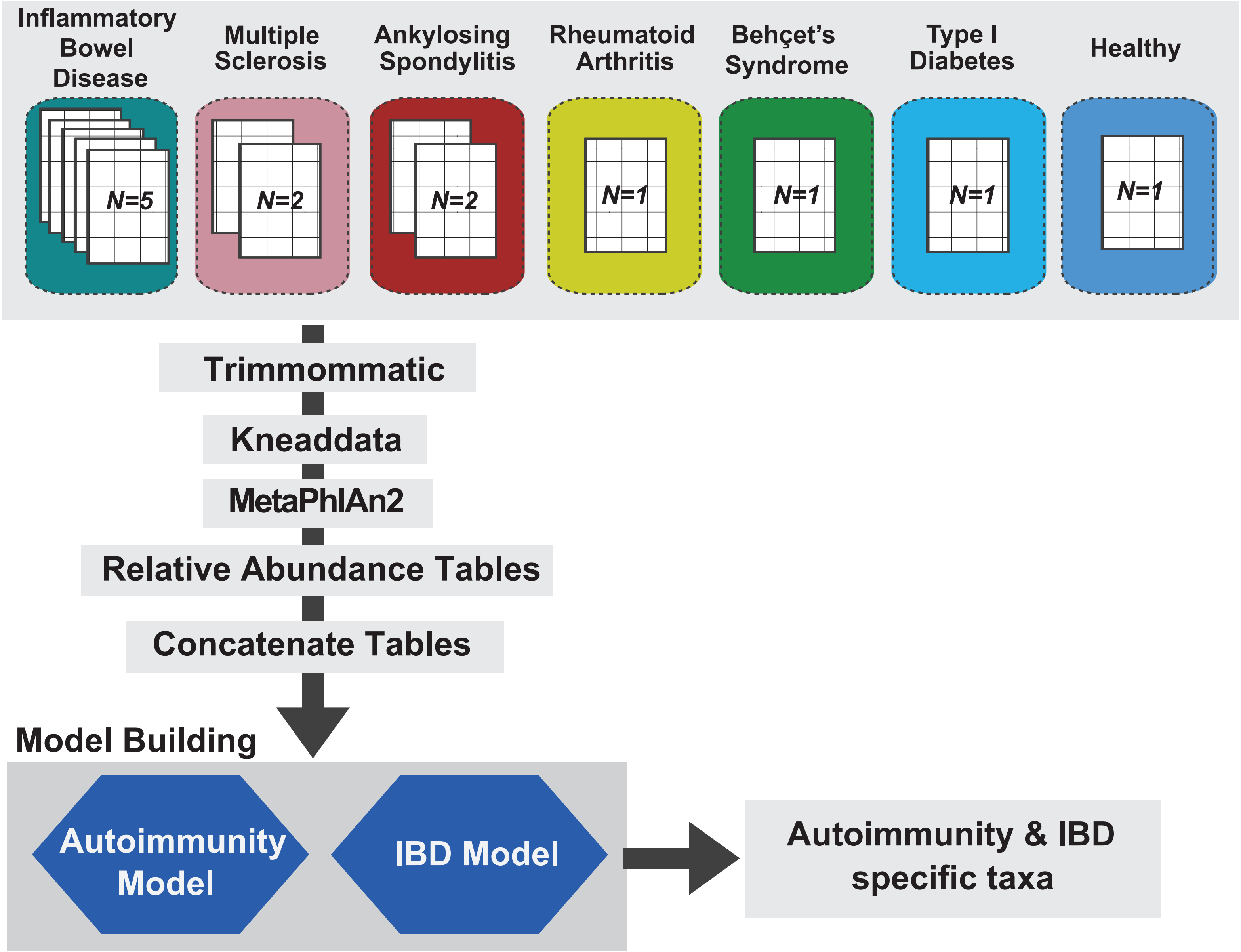

### Supplemental Figure 3

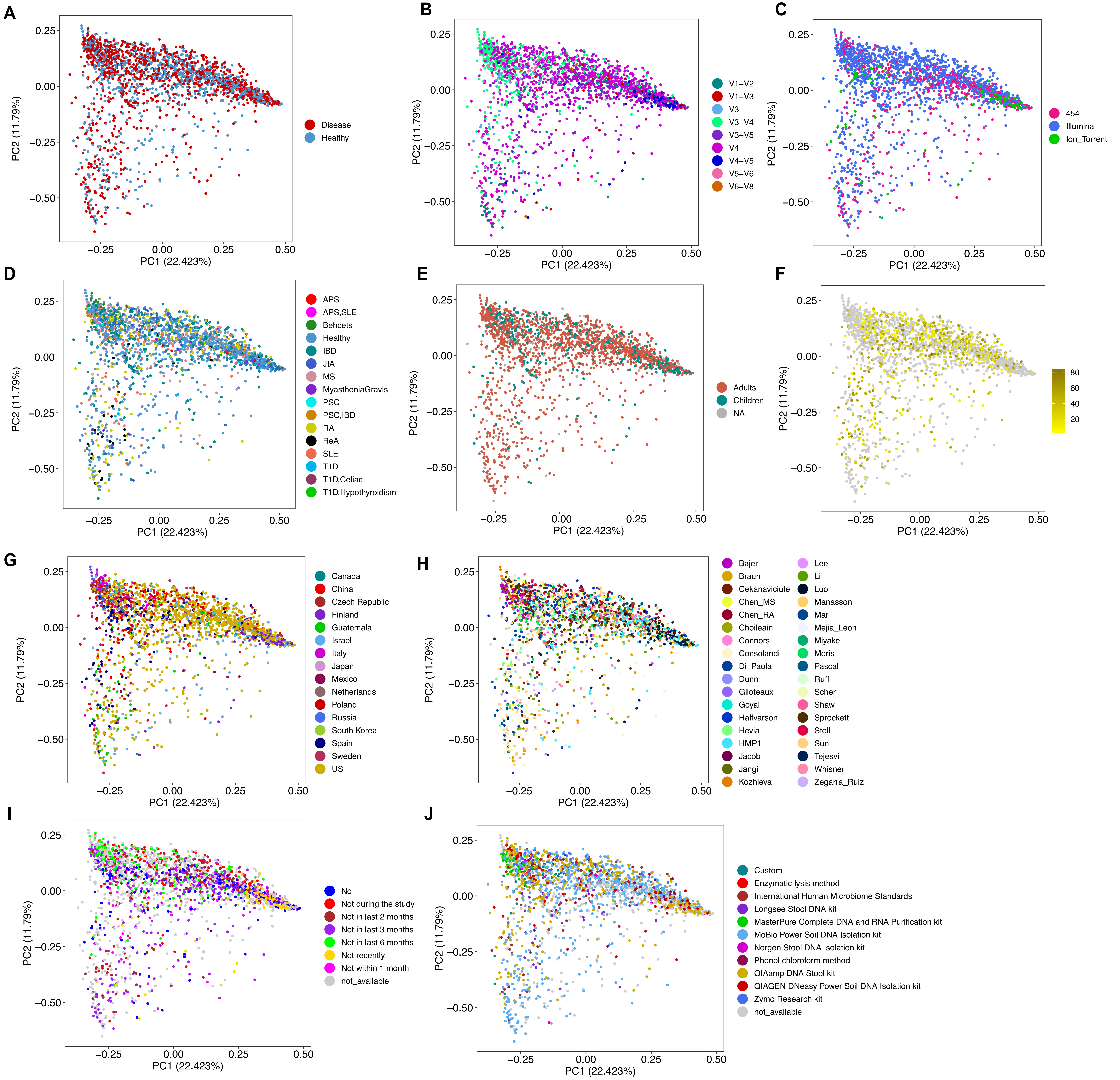

### Supplemental Figure 4

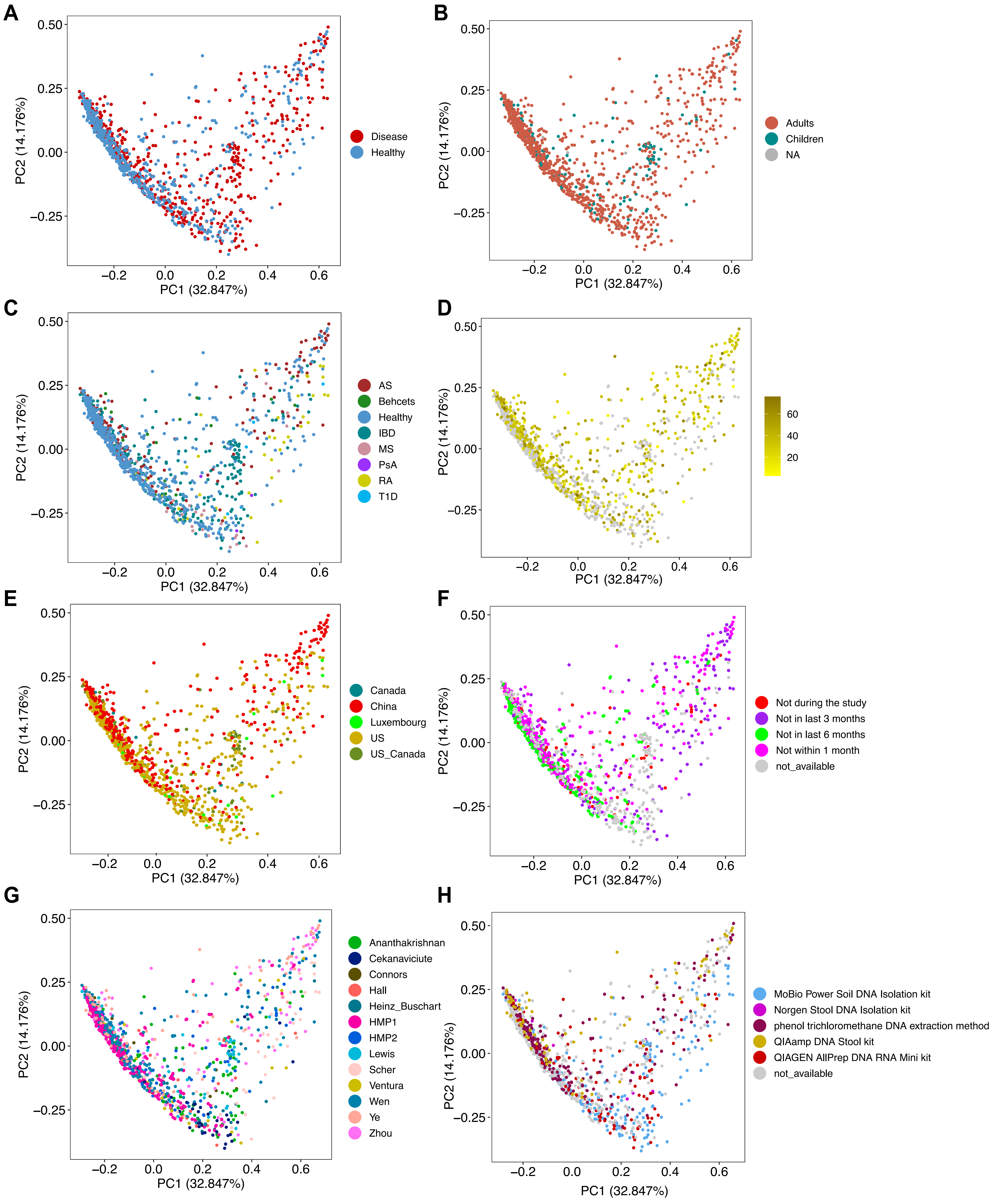

### Supplemental Figure 5

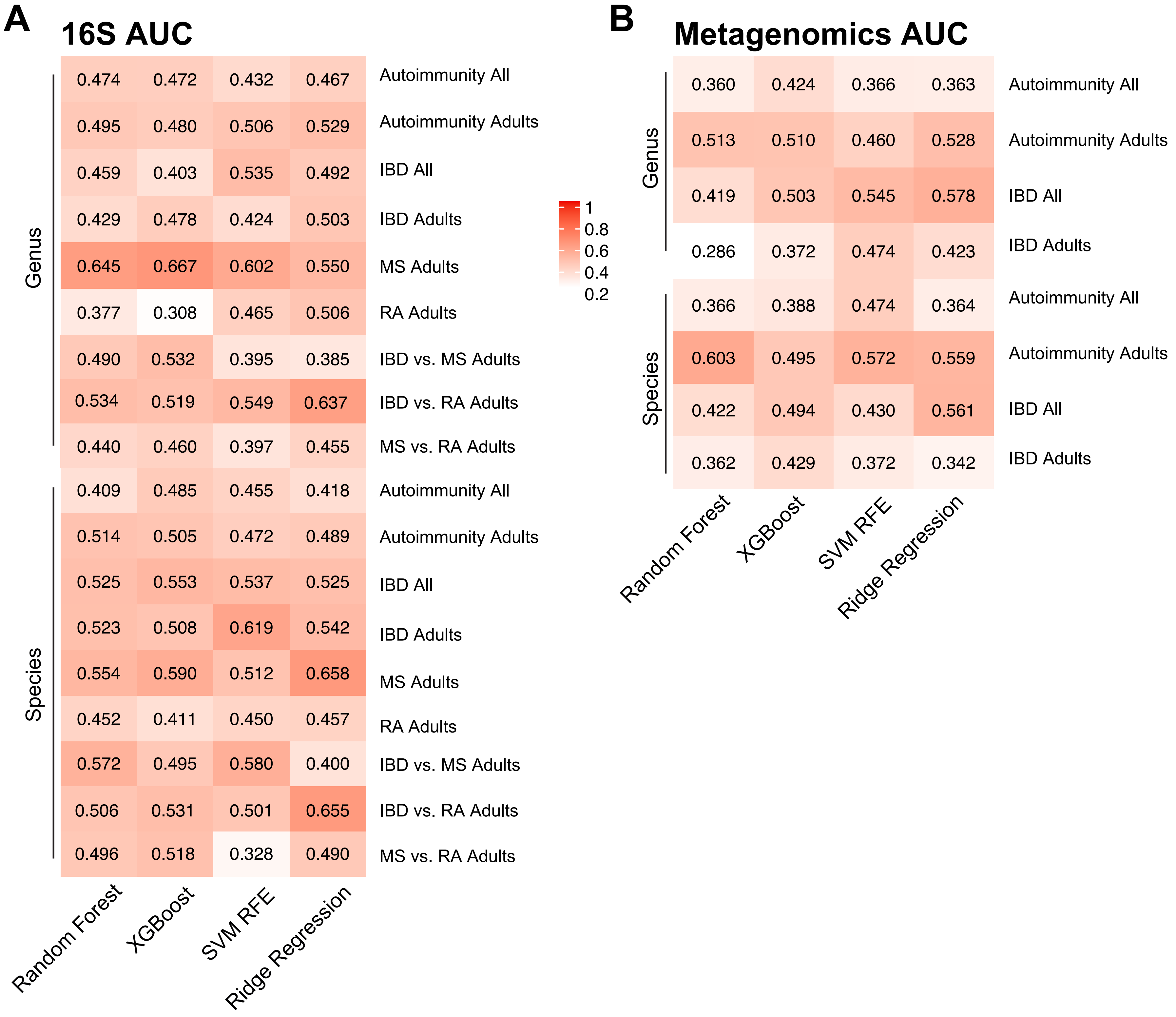

### Supplemental Figure 6AB

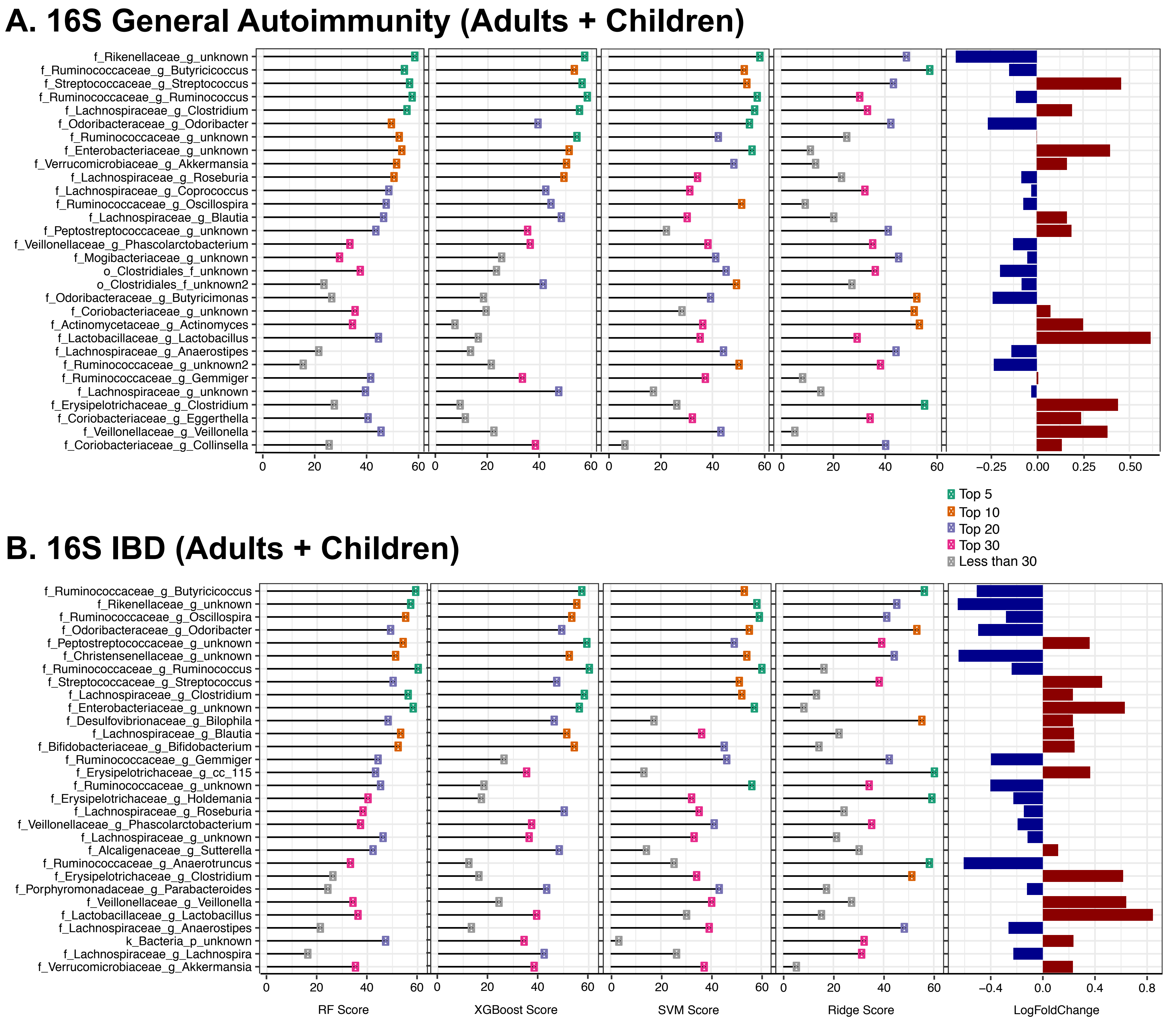

### Supplemental Figure 6CD

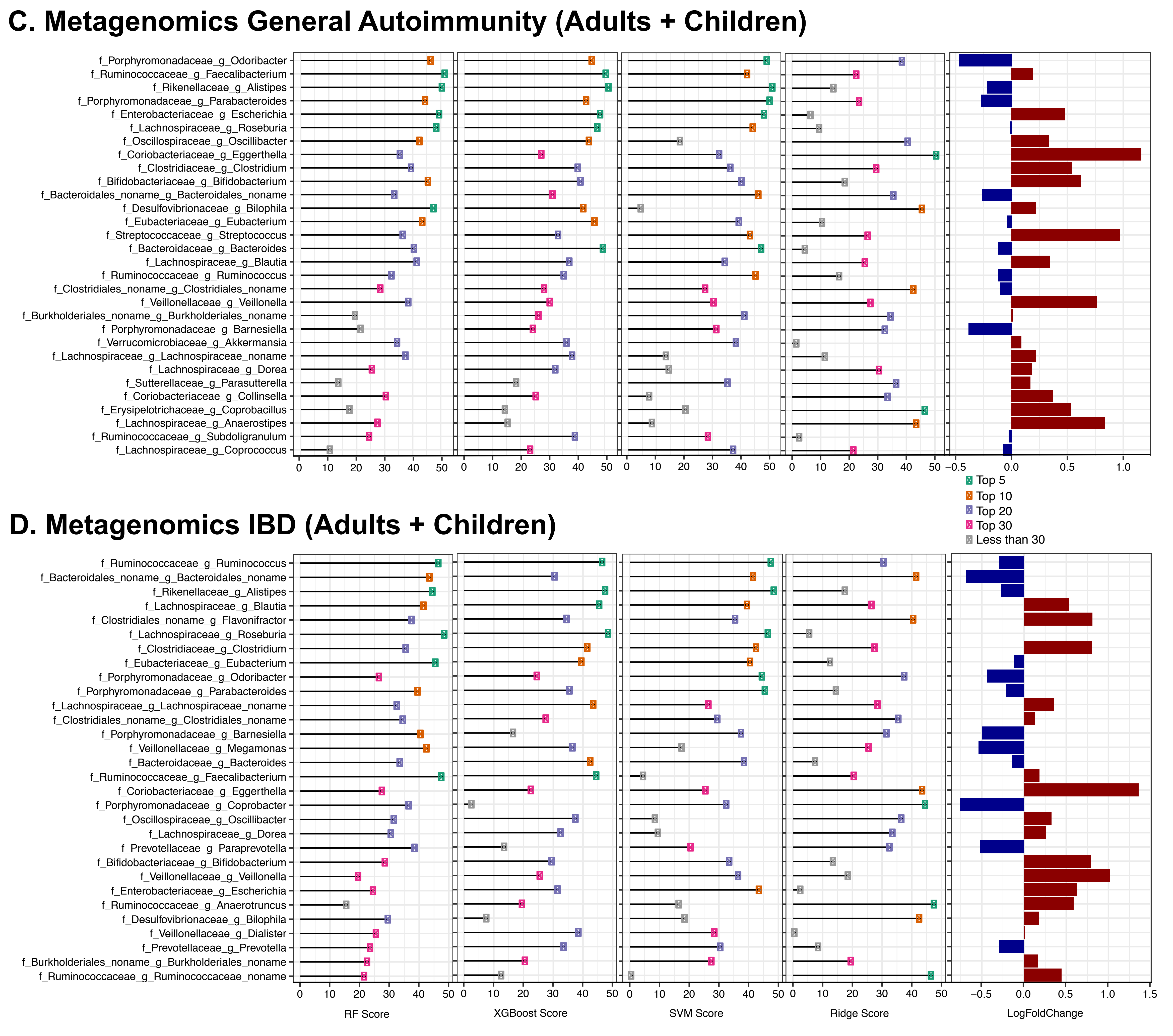

### Supplemental Figure 6EF

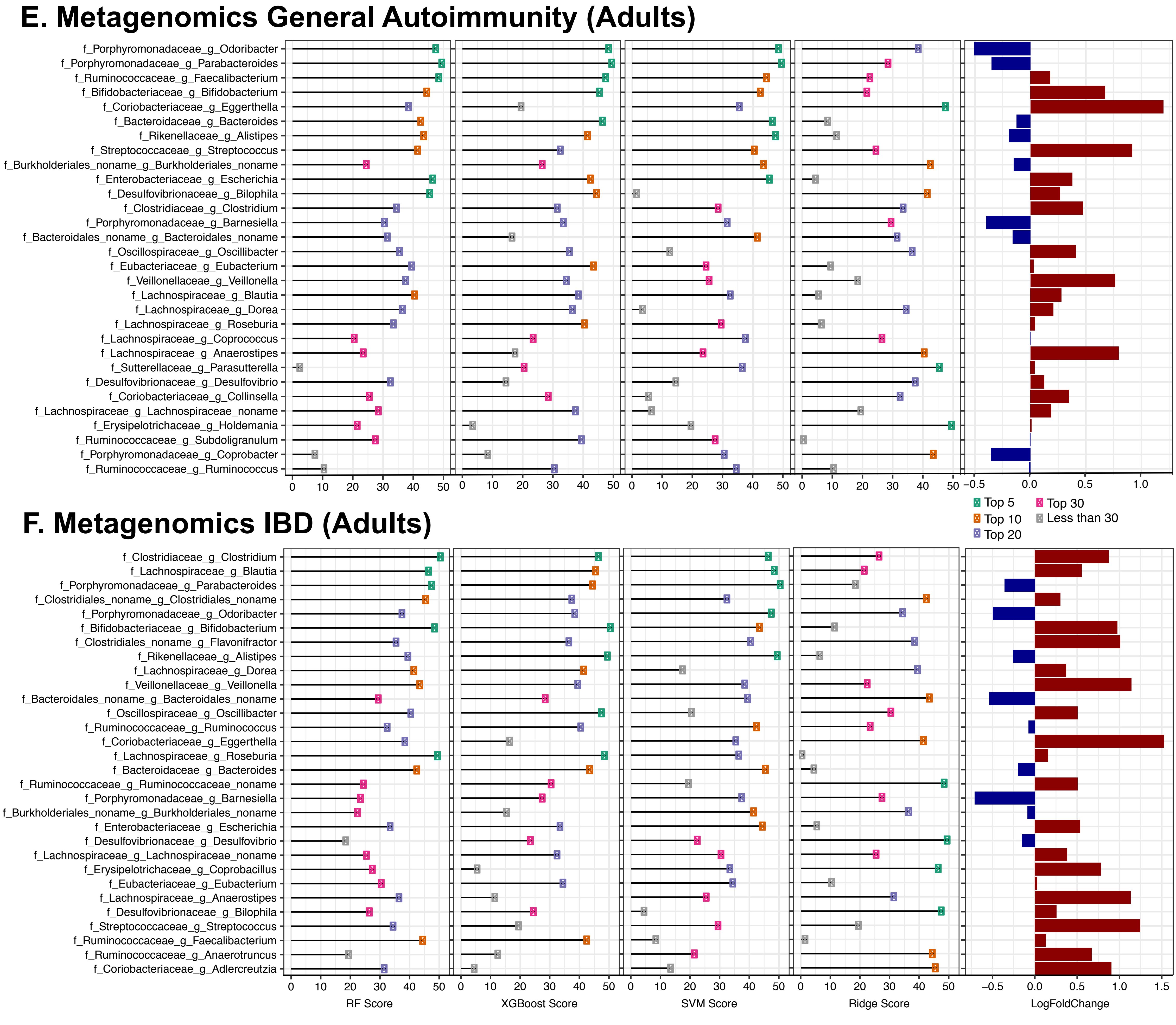

### Supplemental Figure 7

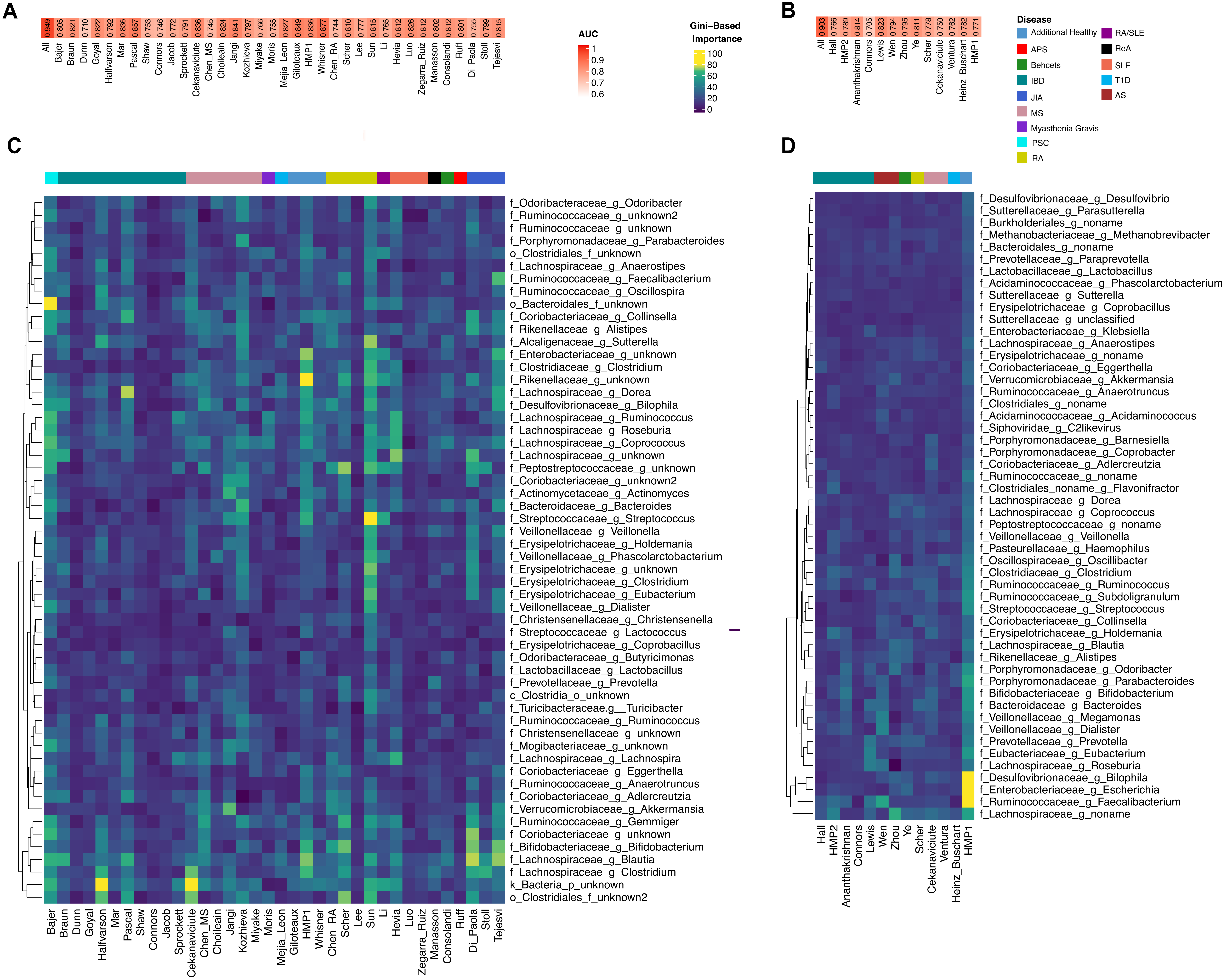

### Supplemental Figure 8

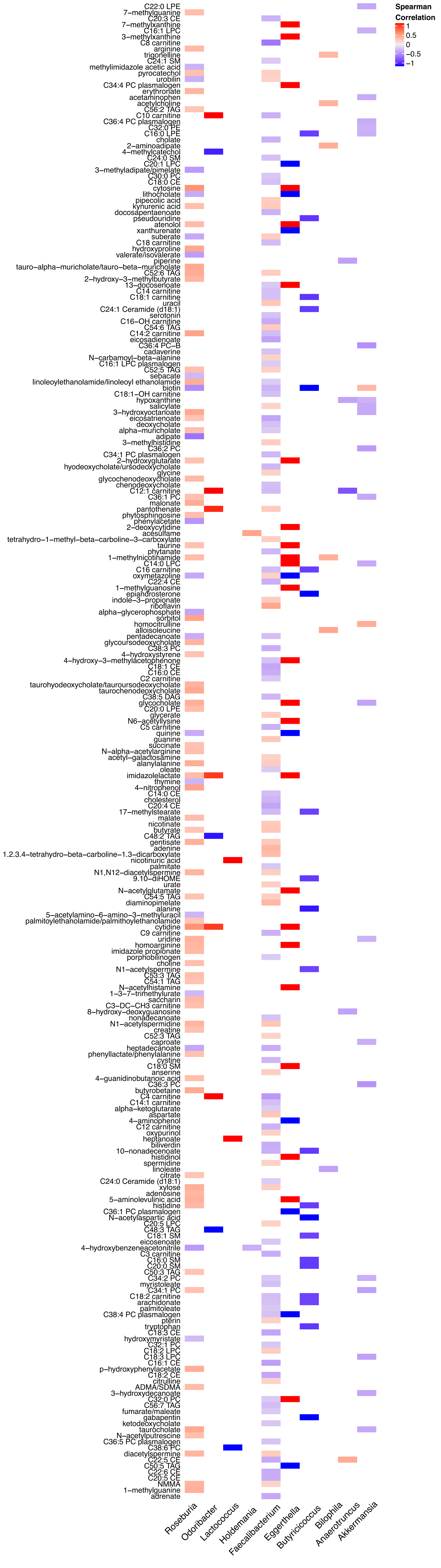
